# Punctuated Aneuploidization of the Budding Yeast Genome

**DOI:** 10.1101/2020.05.04.076935

**Authors:** Lydia R. Heasley, Ruth A. Watson, Juan Lucas Argueso

**Affiliations:** Department of Environmental and Radiological Health Sciences, Colorado State University, Fort Collins-CO, USA

## Abstract

Remarkably complex patterns of aneuploidy have been observed in the genomes of many eukaryotic cell types, ranging from brewing yeasts to tumor cells (*1, 2*). Such aberrant karyotypes are generally thought to take shape progressively over many generations, but evidence also suggests that genomes may undergo faster modes of evolution (*2, 3*). Here, we used diploid *Saccharomyces cerevisiae* cells to investigate the dynamics with which aneuploidies arise. We found that cells selected for the loss of a single chromosome often acquired additional unselected aneuploidies concomitantly. The degrees to which these genomes were altered fell along a spectrum, ranging from simple events affecting just a single chromosome, to systemic events involving many. The striking complexity of karyotypes arising from systemic events, combined with the high frequency at which we detected them, demonstrates that cells can rapidly achieve highly altered genomic configurations during temporally restricted episodes of genomic instability.

## Introduction

Whole chromosome copy number alterations (CCNAs)(*e.g*., aneuploidies) are an important source of phenotypic variation and adaptive potential (*2, 4, 5*). CCNAs usually arise from defects in chromosome segregation (*6*), but, because such errors occur rarely (~10^−6^/cell/division)(*7, 8*), the patterns by which cells accumulate extensive collections of CCNAs remain poorly understood (*2*). Conventional paradigms of genome evolution posit that mutations (*e.g.*, CCNAs) are acquired gradually and independently over many successive generations (*9, 10*). Cancer-centric models have proposed that tumor cells can gain numerous mutations during punctuated and transient bursts of genomic instability (*3, 11–13*), or that they become chronically destabilized and acquire mutations at elevated rates (*i.e*., mutator phenotype)(*14, 15*). Yet, because cancer genome evolution is retrospectively inferred many generations after neoplastic initiation, our understanding of how these mutagenic patterns contribute to the acquisition of CCNAs remains incomplete.

## Results

We used the tractable budding yeast model system to determine the patterns by which CCNAs arise. To recover spontaneously-arising aneuploid clones from populations of diploid cells, we introduced the counter-selectable marker *CAN1* onto the right arm of chromosome V (Chr5R) in the haploid strain JAY291 (*16*). Because the endogenous copy resides on Chr5L, the resulting strain had two copies of *CAN1* on Chr5, one on each arm. We crossed this haploid to the S288c reference strain to form a heterozygous diploid. To select for cells that had lost the JAY291 homolog of Chr5 (jChr5), we grew independent cultures for ≤35 generations in rich media and plated each onto selective media containing canavanine (CAN) (*17*). When we visually inspected CAN-resistant (CAN^R^) colonies, we noted that while the majority had a normal smooth appearance, 1 in ~450 colonies displayed a distinctive rough morphology (Fig. 1A). Previously, we reported that this morphological switch is precipitated by interhomolog mitotic recombination (MR) resulting in loss of the wild type allele of the *ACE2* gene encoded on sChr12R and homozygosis of the mutant *ace2*-*A7* allele on jChr12R (*18*). *ace2-A7* cells fail to separate after cytokinesis and consequently form rough colonies (*18, 19*). In this previous study, rough colonies appeared on non-selective media at a frequency of 1 in ~10,000 colonies and were always caused by MR events spanning *ACE2* on Chr12R (*12, 18*). Rough colonies resulting from whole loss of Chr12 were never observed (0/67 genotyped clones).

**Figure 1.**
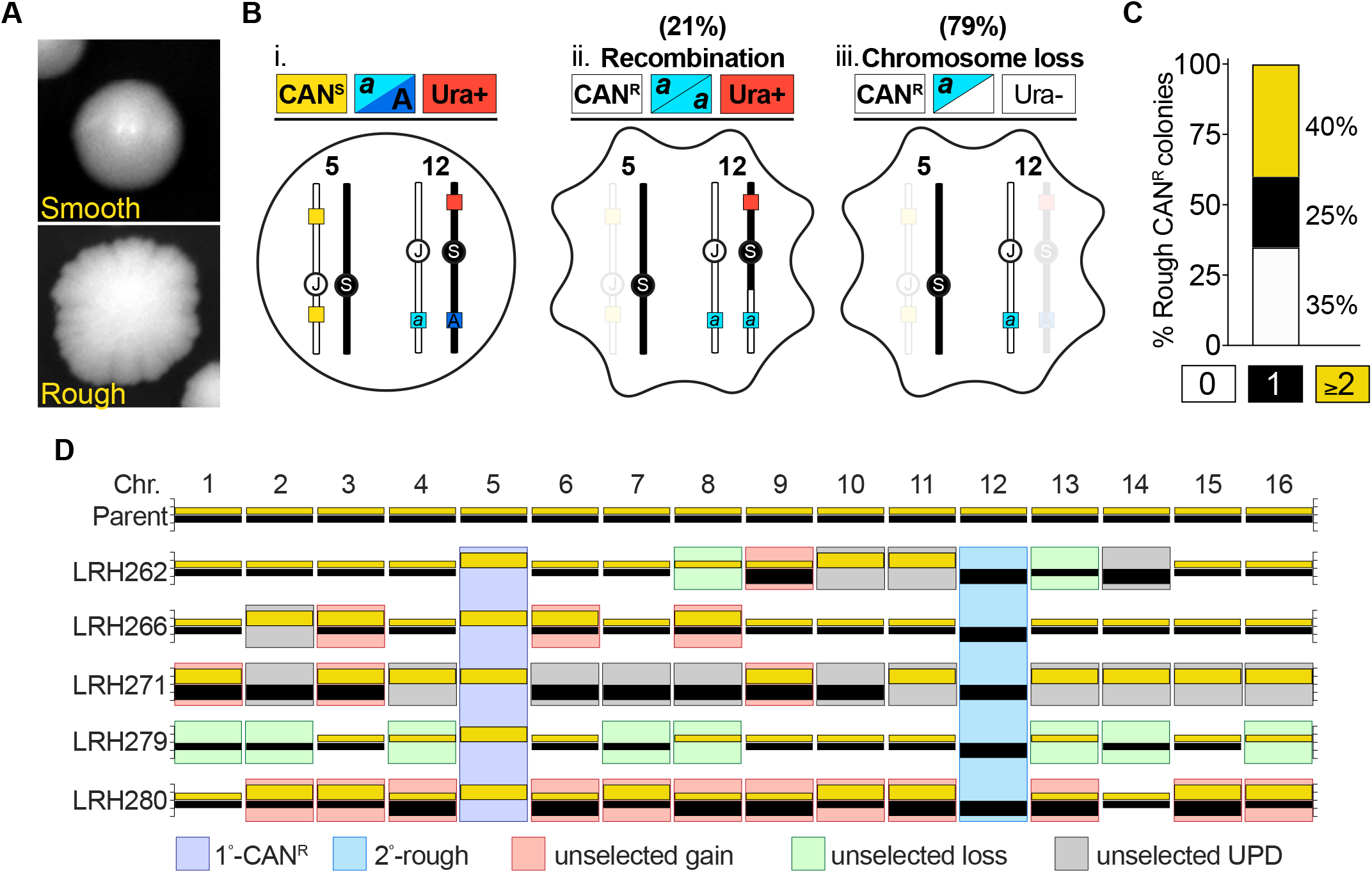
Clones selected for a single CCNA are enriched for additional CCNAs. **(A)** Images of smooth and rough colonies. **(B)** A schematic illustrating the genotypic and phenotypic outcomes of selection for loss of jChr5 and homozygosis of *ace2-A7* on jChr12. jChr5-encoded *CAN1* markers, yellow boxes; jChr12-encoded *ace2-A7* mutation, light blue box; sChr12-encoded *ACE2* allele, dark blue box; sChr12-encoded *URA3* marker, red box. *i.* the parental diploid, *ii.* 21% of rough CAN^R^ colonies were Ura^+^ and homozygous for *ace2-A7* due to MR, *iii.* 79% of rough CAN^R^ were Ura^−^ and hemizygous for *ace2-A7* due to loss of sChr12. **(C)** Percentage of rough CAN^R^ isolates with 0 (white), 1 (black), and ≥2 (yellow) unselected CCNAs. **(D)** Karyotypes of the parent strain and five rough Ura-CAN^R^ isolates. For each chromosome, yellow bars denote the S288c homolog and black bars denote the JAY291 homolog. Colored boxes denote the indicated karyotypic events.

Our finding that rough colonies appeared >22-fold more frequently on CAN selection plates than in non-selective conditions led us to hypothesize that a shared mutational process could have caused the concomitant loss of jChr5 and loss-of-heterozygosity (LOH) on Chr12R. To investigate this, we introduced a *URA3* marker onto sChr12L (Fig. 1B, i). Rough CAN^R^ clones resulting from MR spanning *ACE2* would likely retain this *URA3* marker and grow on media lacking uracil (Ura^+^) (Fig. 1B, ii.), while rough clones caused by loss of the sChr12 homolog would be Ura^−^ (Fig. 1B, iii.). We plated cultures to CAN media, screened CAN^R^ colonies to identify rough clones, and determined the Ura^+/−^ phenotype of each. In contrast to the rough colonies recovered from non-selective conditions (*12, 18*), 79% (41/52) of rough CANR colonies had lost sChr12 in addition to jChr5 (Fig. 1B). Our finding that the selected loss of jChr5 markedly shifted the mutational spectrum of LOH on Chr12R to CCNA was consistent with our above prediction and indicated that clones harboring one aneuploidy were enriched for the presence of additional unselected aneuploidies.

We performed whole-genome sequence (WGS) analysis to comprehensively define the genomic structure of twenty rough CAN^R^ Ura^−^ clones (Table S3). The even distribution of heterozygous sites across the genome of the S288c/JAY291 hybrid enabled us to detect CCNAs of each homolog and changes in overall ploidy. Remarkably, the majority (65%) of the sequenced clones harbored unselected CCNAs of chromosomes other than jChr5 and sChr12 (Fig. 1C, Table S3). Some clones had lost numerous chromosomes (LRH279) while others displayed systemic gains (LRH266 and LRH280)(Fig. 1D). Intriguingly, one clone (LRH271) had acquired CCNAs of every chromosome such that both copies of one homolog had been retained while both copies of the other homolog had been lost, a state known as uniparental disomy (UPD)(*20*). As a result of this UPD-type CCNA, this clone had cumulatively gained and lost 32 homologs and was fully homozygous for either parental haplotype on all chromosomes except Chr1, Chr3, and Chr9, which were tetrasomies (Fig. 1D). The acquisition of such numerous genomic alterations over the limited growth period of ≤35 generations suggested that these clones likely acquired all CCNAs during a temporally restricted episode of chromosomal instability. The homogeneity of WGS read coverage depths observed in the copy number analyses of these clones supported this conclusion. All CCNAs identified within each clonal population were detected at discrete copy numbers; intermediate levels were not observed (data not shown). This demonstrated that CCNAs did not continuously arise during the expansion of the colony, and instead indicated that the instability underlying the formation of these complex genomic alterations was short-lived.

Models of gradual mutation accumulation predict that the rate at which cells independently lose two chromosomes (*2^L^*) should be the multiplicative product of the rates at which each individual chromosome is lost (*1^L^*), referred to here as the *theoretical 2^L^* rate. Our initial results challenged this premise of gradual acquisition and instead suggested that multiple CCNAs could be acquired non-independently. To quantitatively test this gradual model, we constructed a suite of strains in which jChr5 was marked with two copies of *CAN1* and each of several S288c homologs (sChr1, sChr3, sChr9, sChr12) was marked on both arms with copies of *URA3* (Fig. 2A). Plating cultures of these strains to media containing CAN selected for *1^L^* cells that had lost jChr5, and plating to media containing 5-fluoroorotic acid (5-FOA) selected for *1^L^* cells that had lost the *URA3*-marked homolog (*21*). *2^L^* cells that had lost both marked homologs were recovered by plating on media containing both CAN and 5-FOA.

**Figure 2.**
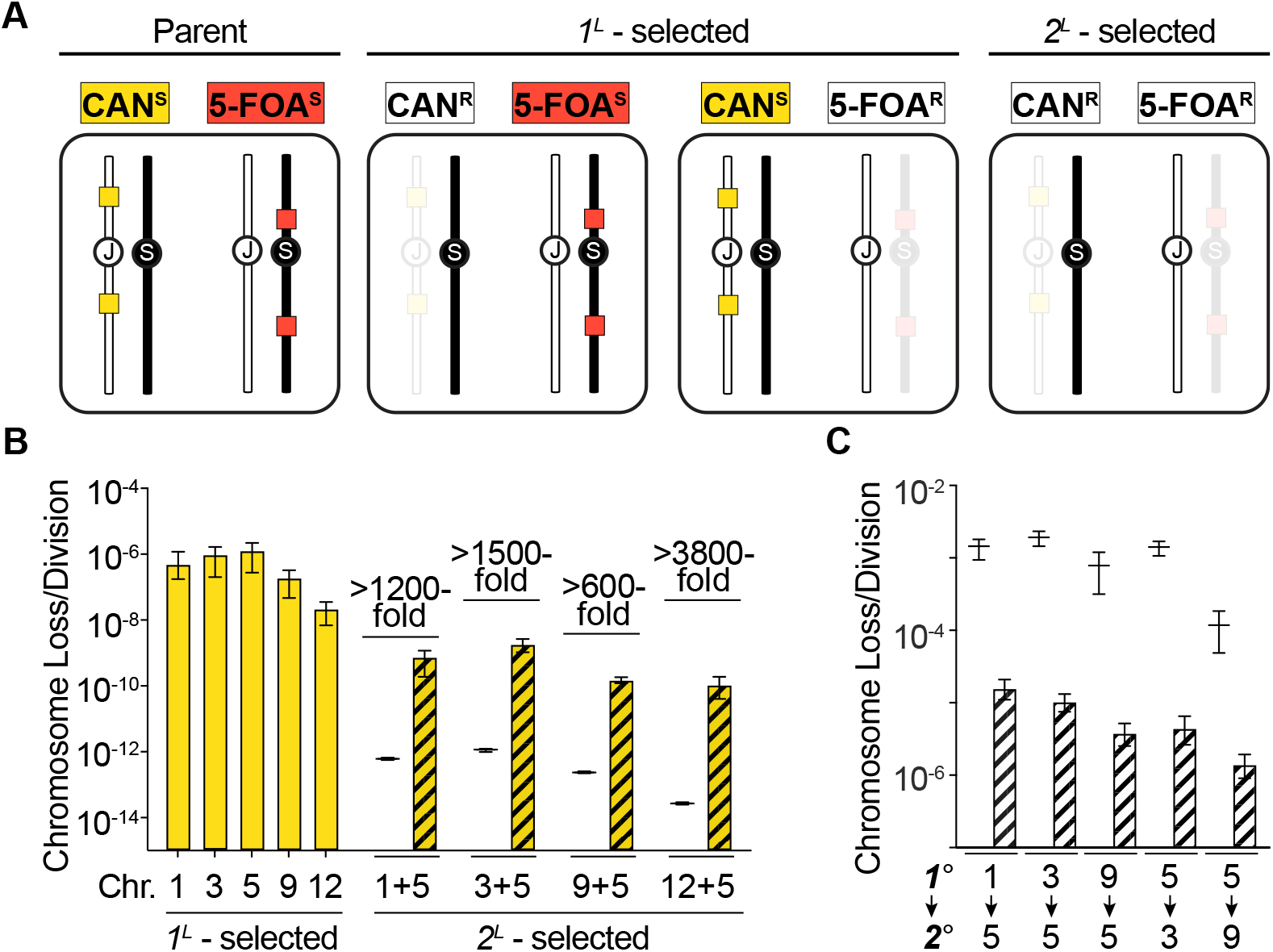
Clones with multiple CCNAs arise more often than predicted by gradual models. **(A)** Schematic illustrating our quantitative CCNA selection approach. jChr5-encoded *CAN1* markers, yellow boxes, S288c homolog-encoded *URA3* markers, red boxes. **(B)** Empirically derived rates of each *1^L^*-selection (yellow) and *2^L^*-selection (yellow striped). Black lines denote *theoretical 2^L^* rates. Fold change between each *theoretical 2^L^* rate and empirically derived *2^L^* rate is noted. **(C)** Empirically derived rates at which cells with a primary existing CCNA (1°) lose a second chromosome (2°)(striped). Black lines denote the theoretical rates at which each 2° CCNA should occur if *2^L^* clones arise by sequential acquisition.

We used fluctuation analysis to determine the rates at which *1^L^* and *2^L^* clones arose in ≤35 generation-cultures (Table S8). Consistent with previous reports (*7, 8*), *1^L^* clones arose at rates of 10^−7^-10^−6^/division (Fig. 2B, yellow bars). Consequently, the *theoretical 2^L^* rates for each pair of aneuploidies were exceedingly low (10^−15^-10^−13^/division; Fig. 2B, black lines). We found that the empirically derived *2^L^* rates were 600-to 3800-fold higher than these *theoretical 2^L^* rates (Fig. 2B, striped bars), demonstrating that *2^L^* clones arise far more frequently than predicted by a gradual model of CCNA acquisition. These results were corroborated by similar experiments in two additional strains (another heterozygous strain S288c/YJM789, and an isogenic strain S288c/S288c; Fig. S1 and Table S8), indicating that the higher-than-expected incidence of *2^L^* clones was a feature common to strains from diverse genetic backgrounds.

In haploids, single aneuploidies can impair chromosomal stability and cause elevated rates of subsequent CCNA acquisition (*22*). We considered the possibility that the *2^L^* clones recovered in our experiments could have resulted from a similar sequential process and tested whether cells aneuploid for a single chromosome exhibited substantially elevated rates of ensuing chromosome loss. If the empirically-derived *2^L^* rates calculated above reflected such a process, then the expected rates at which secondary CCNAs should be acquired would be 1100-fold greater on average (1.2×10^−4^-1.9×10^−3^/division, Fig. 2C, black lines) than the empirically-derived rates of a primary CCNA (Fig. 2B, yellow bars). However, we found that *1^L^* clones (monosomic for sChr1, sChr3, jChr5, or sChr9) lost a second chromosome (jChr5, sChr3, or sChr9) at rates only 2-to 12-fold greater than the euploid parent and far lower than would be expected if *2^L^* clones arose through a process of accelerated sequential accumulation (Table S8). Thus, this effect alone cannot explain the high rates at which *2^L^* clones were recovered in our fluctuation analysis.

We performed WGS analysis of 146 *1^L^* and *2^L^* isolates, as well as fifteen control clones isolated from non-selective conditions. We detected no structural abnormalities in the genomes the control clones. By contrast, and in agreement with our earlier results (Fig. 1), we again observed a remarkable number of *1^L^* and *2^L^* clones containing additional unselected CCNAs (*1^L^:* 39.0%; *2^L^:* 47.9%)(Fig. 3A). Of these unselected CCNAs, each of the sixteen *S. cerevisiae* chromosomes was affected at similar frequencies and we found no evidence that specific CCNAs co-occurred with any particular selected aneuploidy (Fig. 3B). This indicates that unselected CCNAs did not arise subsequently as compensatory suppressors. Additionally, while CCNAs were by far the most prevalent unselected structural genomic alteration, several clones (13/146) had also acquired tracts of LOH resulting from mitotic recombination (Tables S3-S5).

**Figure 3.**
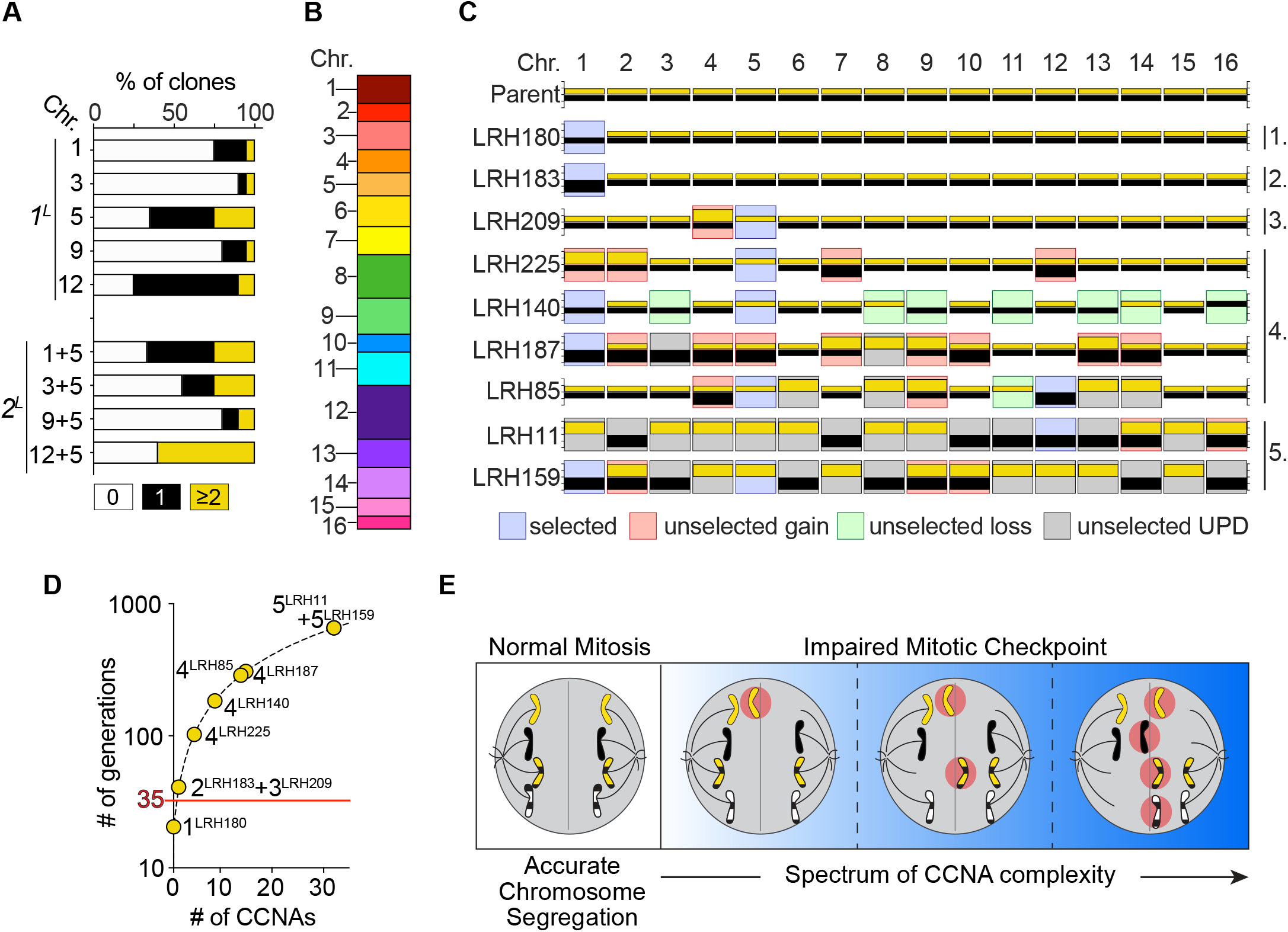
1^L^ and 2^L^ clones display a spectrum of CCNA levels. **(A)** Percentage of *1^L^* and *2^L^* isolates with 0 (white), 1 (black), and ≥2 (yellow) unselected CCNAs. **(B)** Graph depicting the proportion of unselected CCNAs that affected each chromosome. **(C)** Karyotypes of the parent strain and nine clones representing CCNA classes 1-5. Details as in Fig. 1D. **(D)** Plot depicting a model of gradual CCNA accumulation (black dashed line) and the projected number of generations required to generate class 1-5 clones described in (C) (yellow circles). **(E)** A model illustrating how mitosis with impaired checkpoint activity could generate cells with varying numbers of CCNAs. Grey line, division plane. Red circles, mis-segregated chromosomes.

We classified all 146 sequenced clones by the degree to which their genomes had been altered by CCNAs (Fig. 3C). Class 1 clones lost only the selected chromosome(s) and represented 58.2% of the dataset (LRH180, 85/146). The remaining 41.8% of clones contained at least one unselected CCNA (61/146) and were classified as follows: Class 2 clones had additionally gained a second copy of the matched homolog resulting in a UPD-type CCNA (LRH183, 21/61, 34.4%); Class 3 clones harbored one additional CCNA (LRH209, 19/61, 31.1%); Class 4 clones harbored multiple additional CCNAs (LRH225, LRH140, LRH187, LRH85, 19/61, 31.1%); and Class 5 clones harbored UPD-type CCNAs of every homolog (LRH11 and LRH159, 2/61, 3.9%).

We also sequenced the genomes of 86 *1^L^* and *2^L^* isolates derived from the S288c/YJM789 hybrid. Surprisingly, WGS analysis revealed that the parent strain was already trisomic for Chr12 (Fig. S2, Table S5). Despite the this pre-existing CCNA, empirically derived *1^L^* rates for sChr1, sChr3, and yChr5 in this background were comparable to the euploid S288c/JAY291 and S288c/S288c strains (Fig. S1, Table S8). Similar to the clones derived from the S288c/JAY291 hybrid, numerous S288c/YJM789-derived clones contained unselected CCNAs (*1^L^*, 27%; *2^L^*, 40%)(Table S5). Together, CCNA analysis in this background corroborated our above finding that a single pre-existing CCNA, even of a chromosome as large as Chr12, did not substantially perturb genomic stability, nor did it alter the patterns by which derivative clones acquired unselected CCNAs.

We modeled the number of generations required to produce class 1-5 karyotypes shown in Fig. 3C if each CCNA was acquired independently at the average *1^L^* rate of 1.5×10^−6^/division (Fig. 3D, black dashed line). Contrary to our experimental results, this model projected that class 2-5 karyotypes would have required more than 35 generations to develop gradually (41-656 generations)(Fig. 3D, yellow circles). Collectively, the conventional gradual model does not effectively explain the remarkable genomic complexity detected in clones from the datasets above, nor does it account for the frequency at which we recovered such clones. Instead, our results are best explained by a model in which multiple CCNAs are acquired during a transient burst of genomic instability.

Taken together, our results demonstrate the remarkable swiftness with which CCNAs can accumulate to profoundly alter the structure and heterozygosity of a diploid genome. Indeed, cells can and do acquire individual CCNAs independently, indicating that gradual accumulation of CCNAs occurs. But *nearly as often*, cells acquire numerous CCNAs coincidentally. This indicates that a broad spectrum of complex karyotypes can arise during stochastic and short-lived episodes, not as the result of gradualism or chronic genomic instability. Our results in *S. cerevisiae* are directly analogous to recent studies which suggested that it is through this punctuated mode of mutagenesis that cancer cells acquire numerous copy number alterations early in tumorigenesis (*3, 11, 23*). What cellular events might contribute to this process of punctuated copy number evolution (PCNE)(*23*)? Perturbation of many integral cellular processes including DNA damage repair (*24*), replication (*25*), sister chromatid cohesion (*26*), spindle assembly (*27*), and mitotic checkpoint activity (*6*) are known to affect the maintenance and inheritance of chromosomes, and failure of any of these pathways has the potential to affect all chromosomes in a cell equally and simultaneously (*6, 28, 29*). For instance, even a transient failure of the mitotic checkpoint enables a cell to enter anaphase with incorrect chromosome-spindle attachments. Such an erroneous mitosis could produce daughter cells harboring any of the aberrant karyotypic classes described in this study (Fig. 3E)(*6*). Our experimental approach provides a promising model system with which to meticulously define the causal mechanisms of PCNE as well as to assess the phenotypic consequences and adaptive potential of the remarkable karyotypes that can arise from this process.

## Supporting information

Supplementary Table 1. Yeast strains used in this study.

Supplementary Table 2. Plasmids used in this study.

Supplementary Table 3. Sequencing and copy number analysis of rough Ura- CANR S288c/JAY291 clones.

Supplementary Table S4. Sequencing and copy number analysis of 1L and 2L S288c/JAY291 clones.

Supplementary Table S5. Sequencing and copy number analysis of 1L and 2L S288c/YJM789 clones.

Supplementary Table S6. Analysis of the frequency of sequenced clones possessing unselected CCNAs.

Supplementary Table S7. Proportion of unselected CCNAs affecting each chromosome.

Supplementary Table S8. Rates of 1L and 2L chromosome loss calculated using fluctuation analysis.

## Acknowledgements

We are grateful to Eric Alani, Michael McMurray, Dmitry Gordenin and Tom Petes for valuable comments on the manuscript. This study was supported by NIH/NIGMS awards 1K99GM13419301 to LRH and R35GM11978801 to JLA.

## Supplementary methods and associated references

### Strain construction and culture media

All *Saccharomyces cerevisiae* strains used in this study are listed in Table S1 and were derived from the S288c, JAY291 (*16*), or YJM789 (*30*) backgrounds. Plasmids used for PCR-based amplification of selectable markers (*31–33*) are listed in Table S2. Strain construction was performed using standard transformation, crossing, and sporulation procedures. Specific descriptions of the construction of experimental strains are outlined below. To ensure that each strain used in these studies was unable to initiate meiosis and undergo a return-to-growth (RTG) process, we replaced the *IME1* locus on each homolog of Chr10 with *HPHMX* selectable markers. RTG is a process in which diploid yeast cells initiate meiotic programs, introduce Spo11-mediated double strand breaks throughout the genome and then return to vegetative growth (*34*). This process can lead to extensive MR-derived LOH.

### Construction of *CAN1*-marked chromosomes (jChr5, yChr5, sChr5)

A PCR product consisting of *CAN1-KANMX* amplified from genomic DNA was integrated into the *HOM3* locus on Chr5R. Resulting strains had the endogenous *CAN1* gene on Chr5L (31694-33466) and the newly introduced *CAN1-KANMX* cassette on Chr5R (256375-257958).

### Construction of *URA3*-marked chromosomes (sChr1, sChr3, sChr9, sChr12)

The *CORE3* cassette (pJA95), encodes tandem *URA3* genes from *Saccharomyces cerevisiae* (*ScURA3*) and *Kluyveromyces lactis* (*KlURA3*) and a *KANMX* cassette. With the exception of Chr1 (see below), the full *CORE3* marker was introduced on the left arm of each S288c chromosome at the coordinate listed in Table S1. Into an isogenic strain of the opposite mating type, a single *KlURA3* marker was inserted into the right arm of the same chromosome at the coordinate listed in Table S1. The two resulting strains were crossed, sporulated, and spores were dissected to recover a haploid derivative with both the left-arm *CORE3* and right-arm *KlURA3* markers. For construction of *URA3-*encoding sChr1, a *KlURA3* marker was inserted into both the left and right arms.

### Construction of the *TRP1*-marked chromosome (sChr3)

To select for loss of sChr3 in the S288c/YJM789 hybrid, the *TRP1* gene was amplified from genomic DNA and integrated into Chr3L and Chr3R at the coordinates listed in Table S1 in the intermediate strains that were used to make sChr1 (above). These strains were then crossed, sporulated, and spores were dissected to recover a haploid derivative encoding both *TRP1* markers and both *KlURA3* markers. This strain was crossed to JAY2593 to form a heterozygous diploid in which chromosomes sChr1, sChr3, and yChr5 were each marked with counter-selectable markers. Although efficacy of *TRP1* counterselection was strong in the S288c/YJM789 genetic background, we found it to be variable in other genetic backgrounds. For example, we discovered that this selection regime was not effective in an SK1-derived background. Due to the variability of counter-selection efficiency, we used only the *URA3* and *CAN1* counterselection regimes for all experiments in the S288c/JAY291 background.

### Media used to select CCNA clones

Counterselection of *URA3* was performed by plating cells on synthetic complete media (20g/L glucose, 5g/L ammonium sulfate, 1.7g/L yeast nitrogen base without amino acids, 1.4g/L complete drop-out mix, 20g/L bacteriological agar) supplemented with 1g/L 5-Fluoroorotic Acid (5-FOA). Counterselection of *TRP1* was performed by plating cells on synthetic complete media supplemented with .75g/L 5-Fluoroanthranilic Acid (5-FAA). 5-FAA counterselection was only used in plating assays and experiments in the S288c/YJM789 background. Counterselection against *CAN1* was performed by plating cells on synthetic media lacking arginine (20g/L glucose, 5g/L ammonium sulfate, 1.7g/L yeast nitrogen base without amino acids, 1.4g/L arginine dropout mix, 20g/L bacteriological agar) supplemented with 0.06g/L canavanine sulfate (CAN). Selection of *2^L^* clones was performed by plating cells to appropriate media supplemented with 1g/L 5-FOA and 0.06g/L CAN, 1g/L 5-FOA and 0.75g/L 5-FAA (S288c/YJM789 only), or 0.75g/L 5-FAA and 0.06g/L CAN (S288c/YJM789 only). Because most S288c chromosomes in the isogenic experiments were marked with *URA3* cassettes, selection of the *2^L^* combinations sChr1/sChr3, sChr1/sChr9, and sChr1/sChr12 was conducted by plating cells to media supplemented with 1g/L 5-FOA.

### Rough Colony Screening and Analysis

Diploid yeast cells of the strain JAY2775 were streaked on solid YPD media and incubated at 30°C for 32 hours to allow single colonies to grow. Single colonies were each inoculated into 5 or 7mL liquid YPD cultures and incubated at 30°C for another 24 hours on a rotating drum. Each culture was then diluted appropriately, plated onto CAN-supplemented media, and incubated at 30°C for 4 days. Plates were then visually screened for the presence of rough colonies. Rough colonies were isolated with a sterile toothpick and streaked onto both CAN-supplemented media (to preserve a stock) and uracil-dropout media (20g/L glucose, 5g/L ammonium sulfate, 1.7g/L yeast nitrogen base without amino acids, 1.4g/L uracil drop-out mix, 20g/L bacteriological agar). Plates were incubated at 30°C for 24 hours. After 24 hours, each clone was assessed for its ability to grow on uracil-dropout media.

### Genome sequencing and analysis

The genomes of 276 unselected, *1^L^*, and *2^L^* clones from either the S288c/JAY291 or S288c/YJM789 hybrid backgrounds were sequenced using Illumina short read whole genome sequencing. Genomic DNA from each clone was isolated using the Yeastar Genomic DNA kit from Zymo Research. Pooled, barcoded libraries of 96 individual genomes were generated using Seqwell plexWell-96 kits. Each 96-sample library was sequenced on a single Illumina HiSeq lane. Using CLC Genomics Workbench software (Qiagen), the following processing pipeline was utilized to analyze each sequenced genome: Illumina reads for each genome were imported into CLC and mapped to the most recent release of the yeast reference genome (R64-2-1, yeastgenome.org). Each resulting read mapping file was then imported into the Nexus Copy Number software (Biodiscovery). Each file was subjected to copy number and single nucleotide polymorphism (SNP) variant analysis to identify the copy number of each chromosome (relative to the diploid parent) and heterozygosity at >20,000 individual sites distributed across the genome. From this, we identified the following structural variations: whole chromosome gains/losses, segmental duplications/deletions, and tracts of loss-of-heterozygosity (LOH). LOH breakpoints identified in Nexus were confirmed manually in CLC (Tables S3-S5).

Two different approaches were used to define CCNAs, and the analysis of each sequenced dataset are present in Table S6: 1) the 16-chromosome pairs method; aneuploidy was defined as the deviation of overall ploidy away from 2n. Using this method, uniparental disomy was not scored as an aneuploidy, despite loss of one homolog and gain of the other homolog, 2) the 32-homologs method; aneuploidy was defined as the deviation in copy number of each individual homolog away from 1n. Using this method, UPDs were scored as two CCNAs. Graphs in Figs. 1C, 3A, and S2A depict the results from the 32-homologs method of analysis. Results from both the 16-chromosome pairs and 32-homologs analyses for each sequenced dataset are presented in Table S6.

Graphs in Figs. 3B and S2B depict the proportion of total unselected aneuploidies that affected each yeast chromosome. To determine if there was a bias towards any chromosome in terms of gains/losses, we used the chi square goodness of fit test to compare the distribution of observed frequency of CCNA for each chromosome to the test distribution of expected null rates of 6.25% per chromosome (100% divided by 16 chromosomes). From this test, we calculated a p-value of 0.109, which indicated that there was no significant difference between each chromosome. Because we found no evidence of biases favoring specific chromosomes, we pooled the total number of unselected aneuploidies in the complete S288c/JAY291 or S288c/YJM789 dataset regardless of primary selection (*e.g.,* selection for loss of sChr1). These data are presented in Table S7.

### Data Availability

Sequence files for each clone in this study are available through NCBI (SRA) SUB7254181. All strains and raw data presented in this study will be shared upon request.

### Quantitative Chromosome Loss Assays

Cultures of S288c/JAY291 diploid strains were prepared from single colonies in a manner identical to that used to select for rough CAN^R^ clones (see above). Each culture was serially diluted and plated onto YPD (non-selective), 5-FOA- and CAN-supplemented medias (*1^L^* selection), and 5-FOA+CAN-supplemented media (*2^L^* selection). For the experiments using the S288c/YJM789 diploid strains, cultures were also plated onto 5-FAA-supplemented media (sChr3 *1^L^* selection), and onto 5-FOA+5-FAA- and CAN+5-FAA-supplemented media (*2^L^* selection). Colonies on non-selective and *1^L^*-selected plates were counted after 4 days of growth. Colonies on *2^L^*-selected plates were counted after 6 days of growth. Colony count data were used to calculate rates and 95% confidence intervals of chromosome loss using Flucalc, a MSS-MLE (Ma-Sandri-Sarkar Maximum Likelihood Estimator) calculator for Luria-Delbrück fluctuation analysis (flucalc.ase.tufts.edu)(*35*). To determine the *theoretical* rates at which *2^L^* clones should arise if each chromosome was lost independently, the multiplicative product of both observed *1^L^* rates (and corresponding 95% confidence intervals) was calculated as follows: theoretical rate *2^L(ChrA+ChrB)^* = empirically-derived rate *1^L(ChrA)^* x empirically-derived rate *1^L(ChrB^*^)^. The following rationale was used to calculate the theoretical rates of sequential secondary CCNA acquisition depicted in Fig. 2C (black lines). Using empirically-derived *1^L^* and *2^L^* rates (Fig. 2B and Table S8), we calculated the rate at which a secondary chromosome (ChrB) would be expected to be lost following loss of a primary chromosome (ChrA) if due to sequential process: theoretical sequential rate *1^L(ChrB)^* = empirically-derived rate *2^L(ChrA+ChrB)^* / empirically-derived rate *1^L(ChrA)^*. All empirically derived and *theoretical* rates, 95%-confidence intervals, and number of cultures used to calculate each rate are listed in Table S8.

### Modeling gradual acquisition of CCNAs

We modeled the generations associated with the gradual acquisition of CCNAs using the equation #gen=Log2((1.5×10^6^)^#A^) in which #gen equals the number of generations, #A equals number of CCNAs, and 1.5×10^6^ defines a representative and constant rate of chromosome loss.

---

**Figure S1.**
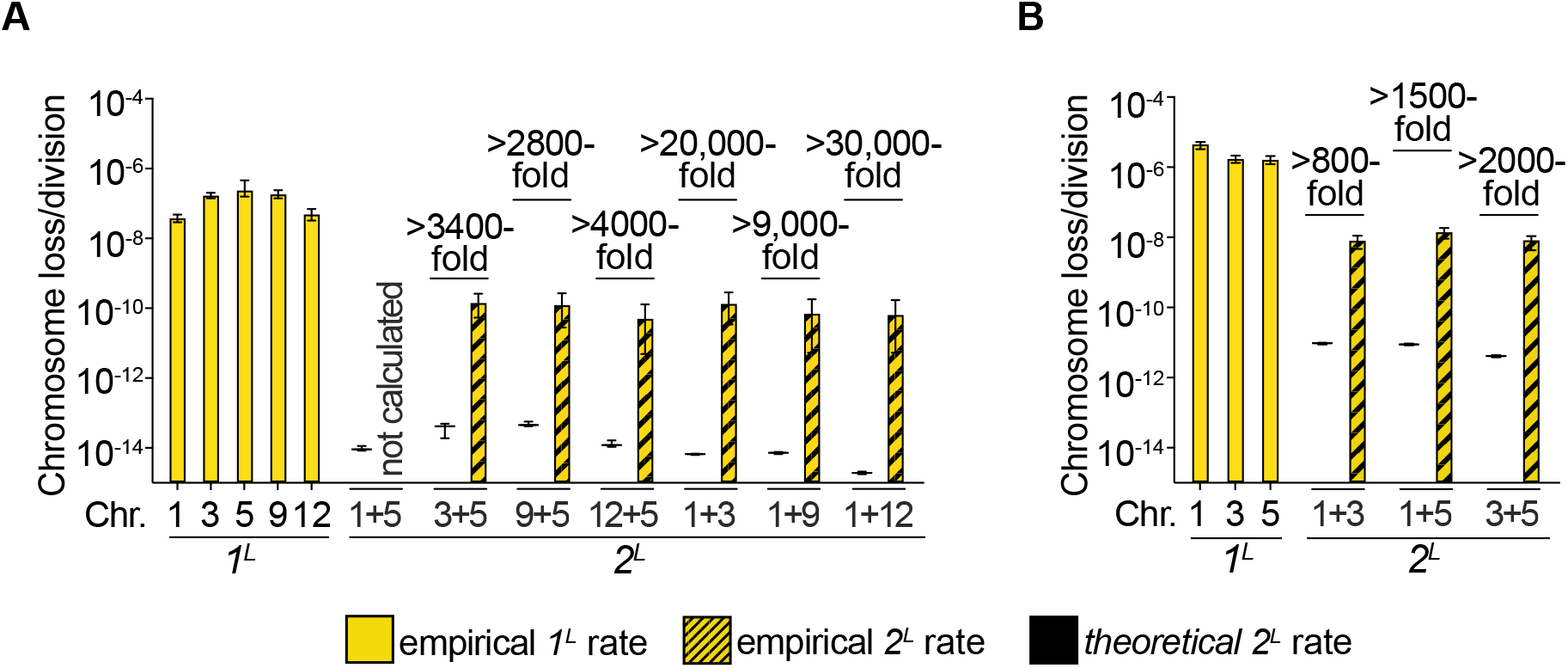
*1^L^* and *2^L^* rate analysis in two diverged genetic backgrounds. **(A)** Empirically derived rates of chromosome loss for each *1^L^*-selection (yellow) and *2^L^*-selection (yellow striped) in an isogenic S288c/S288c background. Black lines denote *theoretical 2^L^* rate predictions. Fold change between *theoretical 2^L^* rates and empirically derived *2^L^* rates (black lines vs. yellow striped) are noted. **(B)** Empirically derived rates of chromosome loss for each *1^L^*-selection (yellow) and *2^L^*-selection (yellow striped) in the hybrid S288c/YJM789 background. Black lines denote *theoretical 2^L^* rate predictions. Fold change between *theoretical 2^L^* rates and empirically derived *2^L^* rates (black lines vs. yellow striped) are noted.

**Figure S2.**
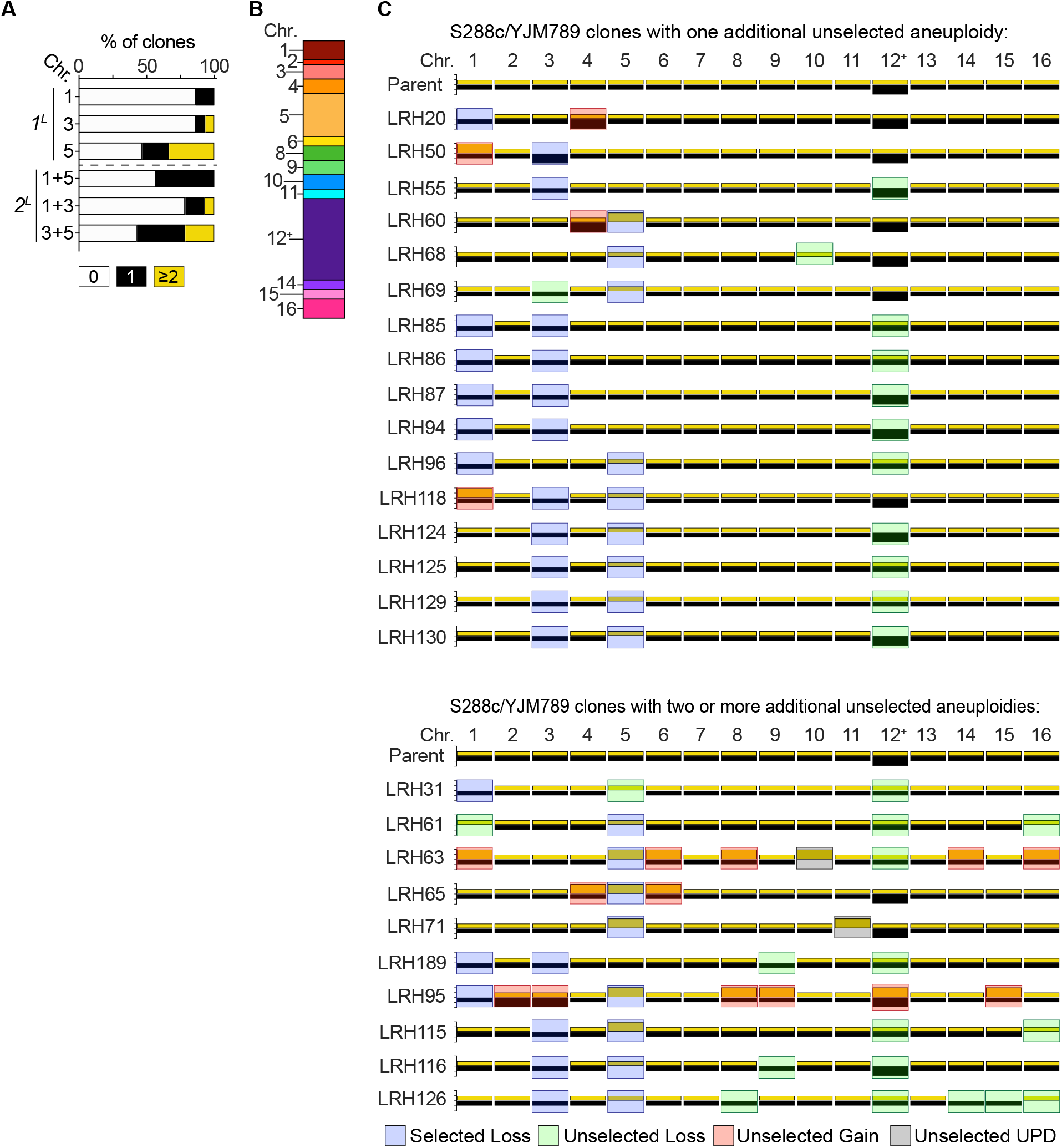
Genomic analysis of S288c/YJM789 *1^L^* and *2^L^* clones. **(A)** Percentage of *1^L^* and *2^L^* isolates with 0 (white), 1 (black), and ≥2 (yellow) unselected CCNAs. **(B)** Graph depicting the proportion of unselected CCNAs affecting each chromosome. Note that cells were trisomic for Chr12 (12^+^). **(C)** Karyotypes of the parent strain and all clones containing ≥1 unselected CCNA. For each chromosome, yellow bars denote the S288c homolog and black bars denote the YJM789 homolog. yChr12 is present at two copies. Colored boxes represent denoted karyotypic events.

**Supplementary Table 1.**
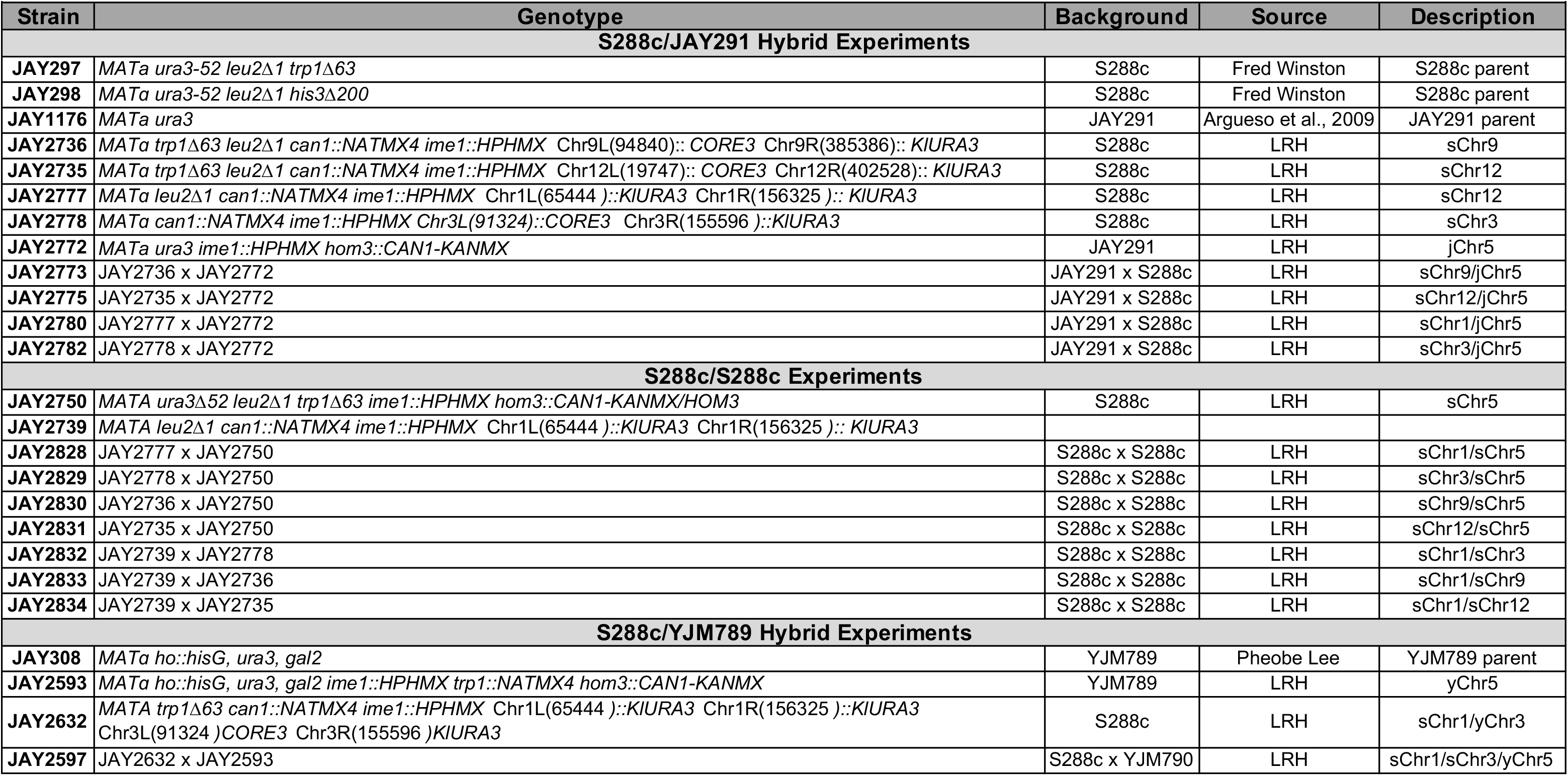
Yeast Strains Used in This Study

**Supplementary Table 2.**
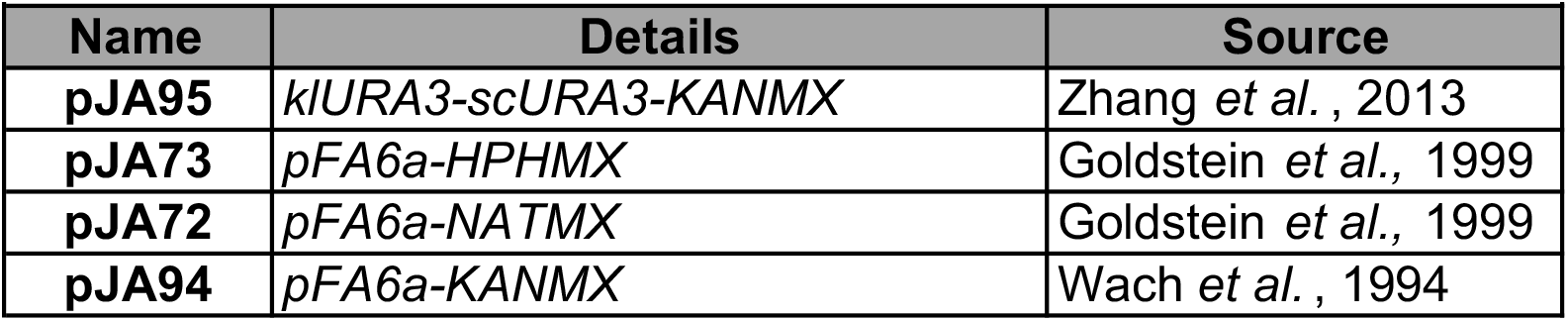
Plasmids used in this study

**Supplementary Table 3.**
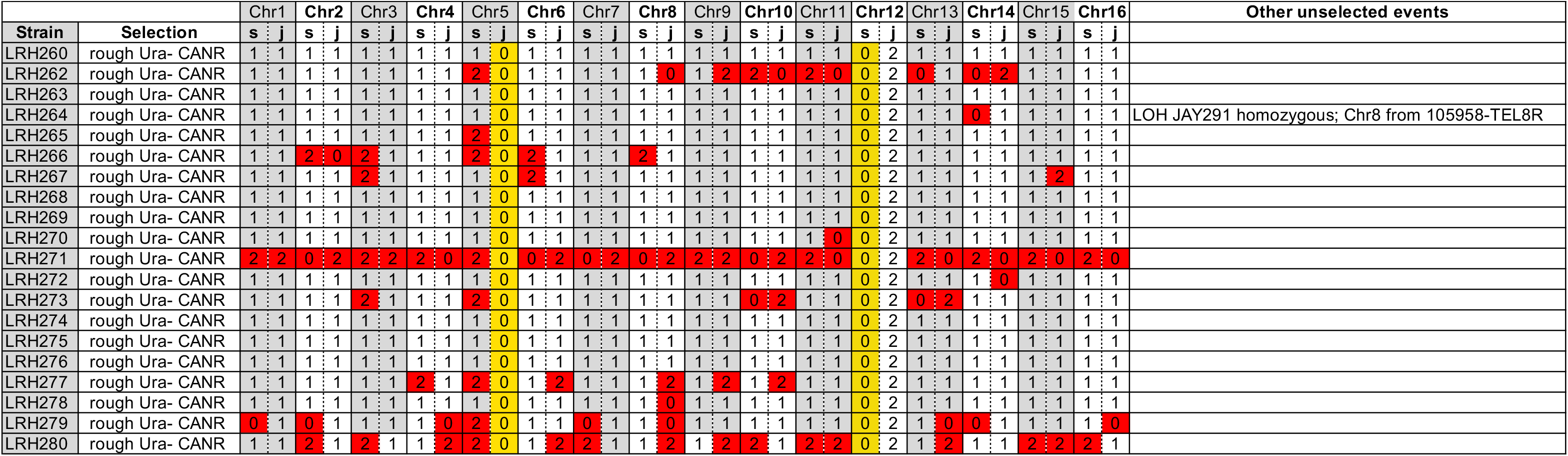
Sequencing and copy number analysis of rough Ura-CANR S288c/JAY291 clones. Columns: s=S288c, j=JAY291. Selected CCNAs (jChr5 and sChr12) are highlighted in yellow. Unselected CCNAs are highlighted in red. Note: All CCNAs of sChr12 were UPD-type.

**Supplementary Table 4.**
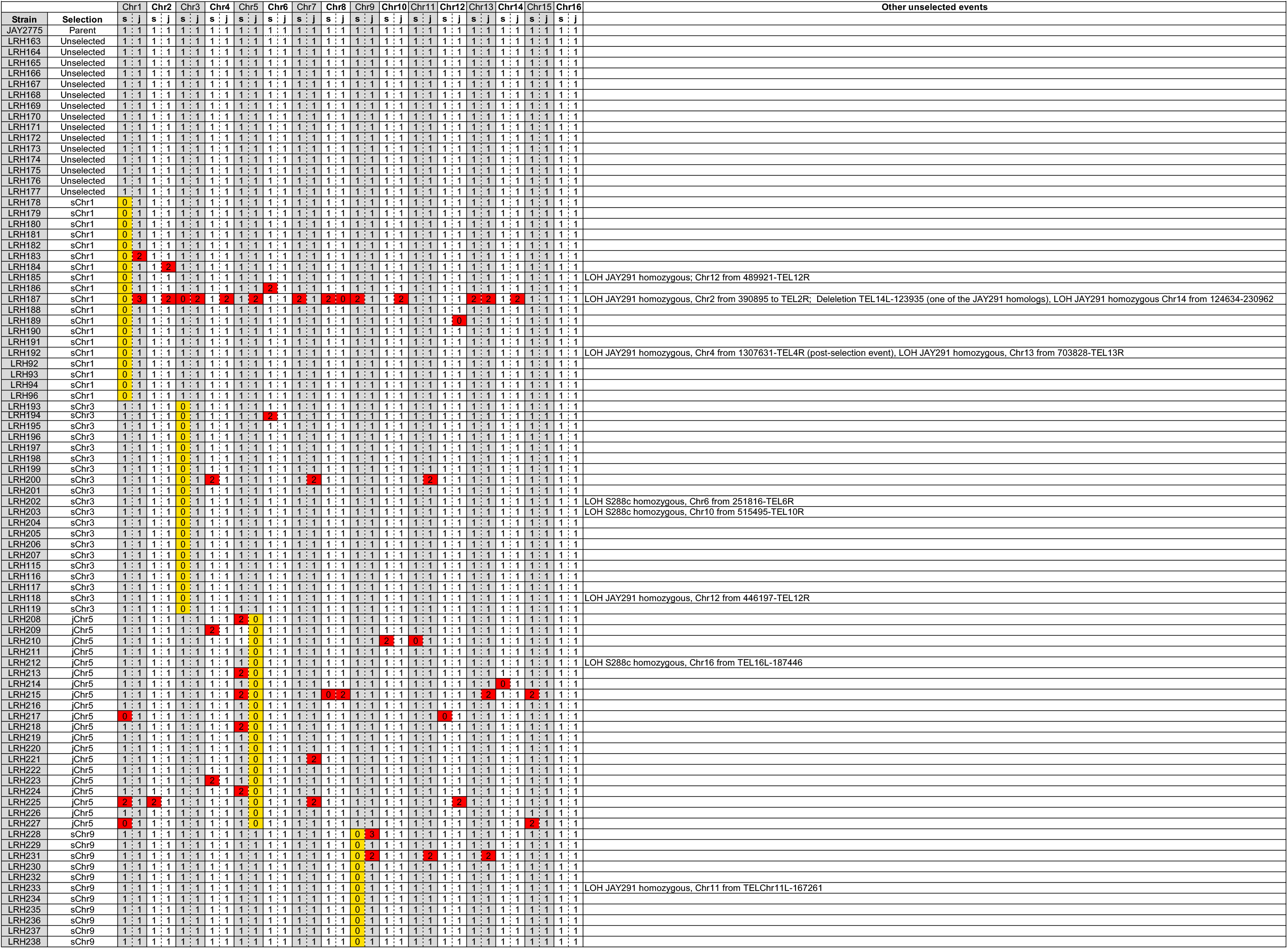
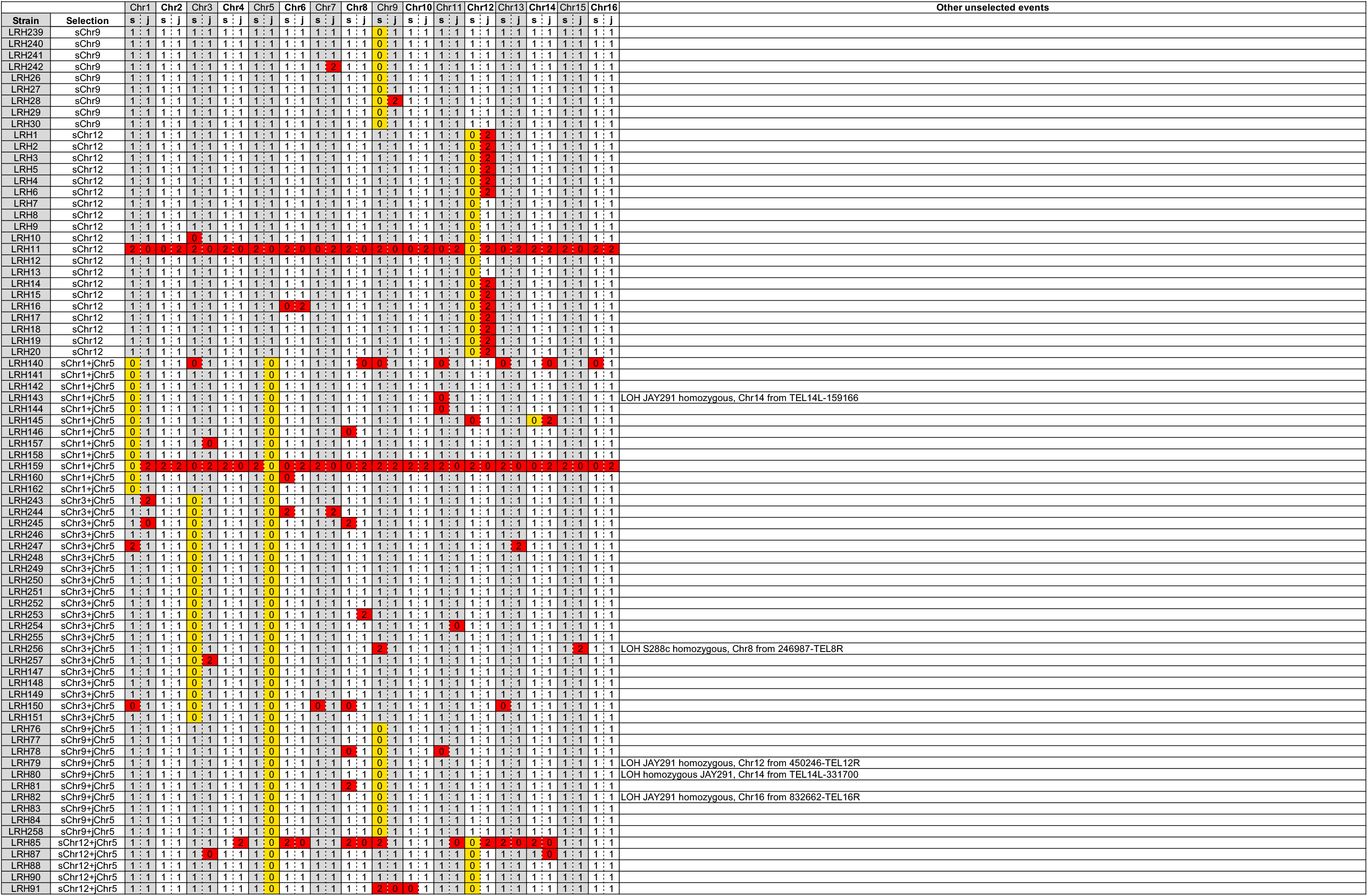
Sequencing and copy number analysis of *1L* and *2L* S288c/JAY291 clones. Columns: s=S288c, j=JAY291. Selected CCNAs are highlighted in yellow. Unselected CCNAs are highlighted in red.

**Supplementary Table 5.**
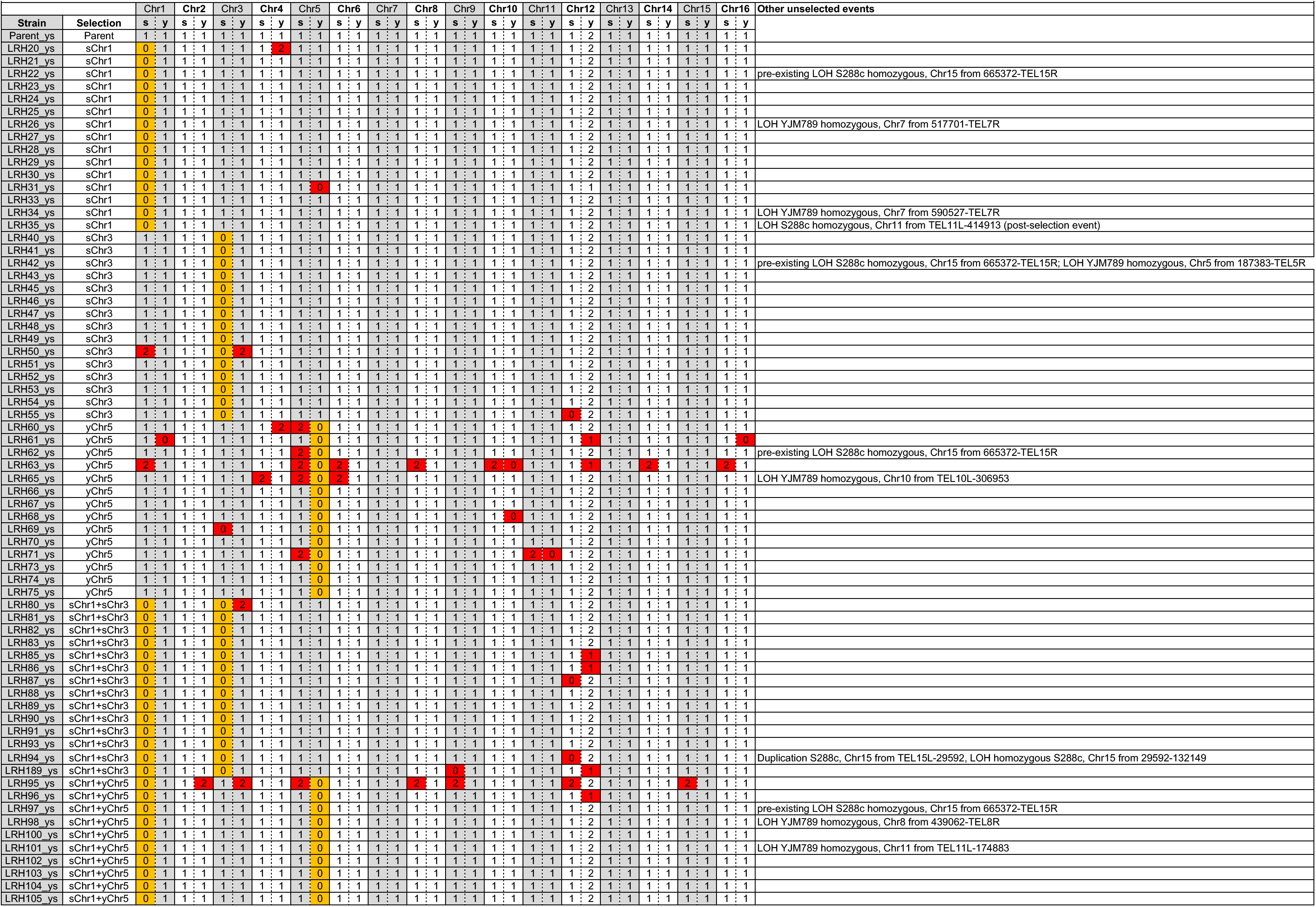
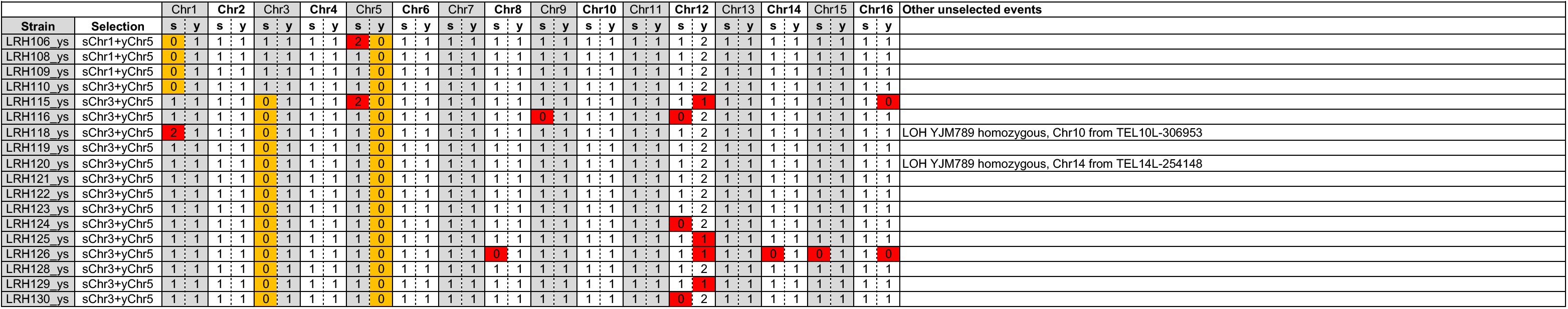
Sequencing and copy number analysis of *1L* and *2L* S288c/YJM789 clones. Note: Diploid parent is trisomic for Chr12 (2 copies of YJM789 Chr12). Columns: s=S288c, y=YJM789. Selected CCNAs are highlighted in yellow. Unselected CCNAs are highlighted in red.

**Supplementary Table 6.**
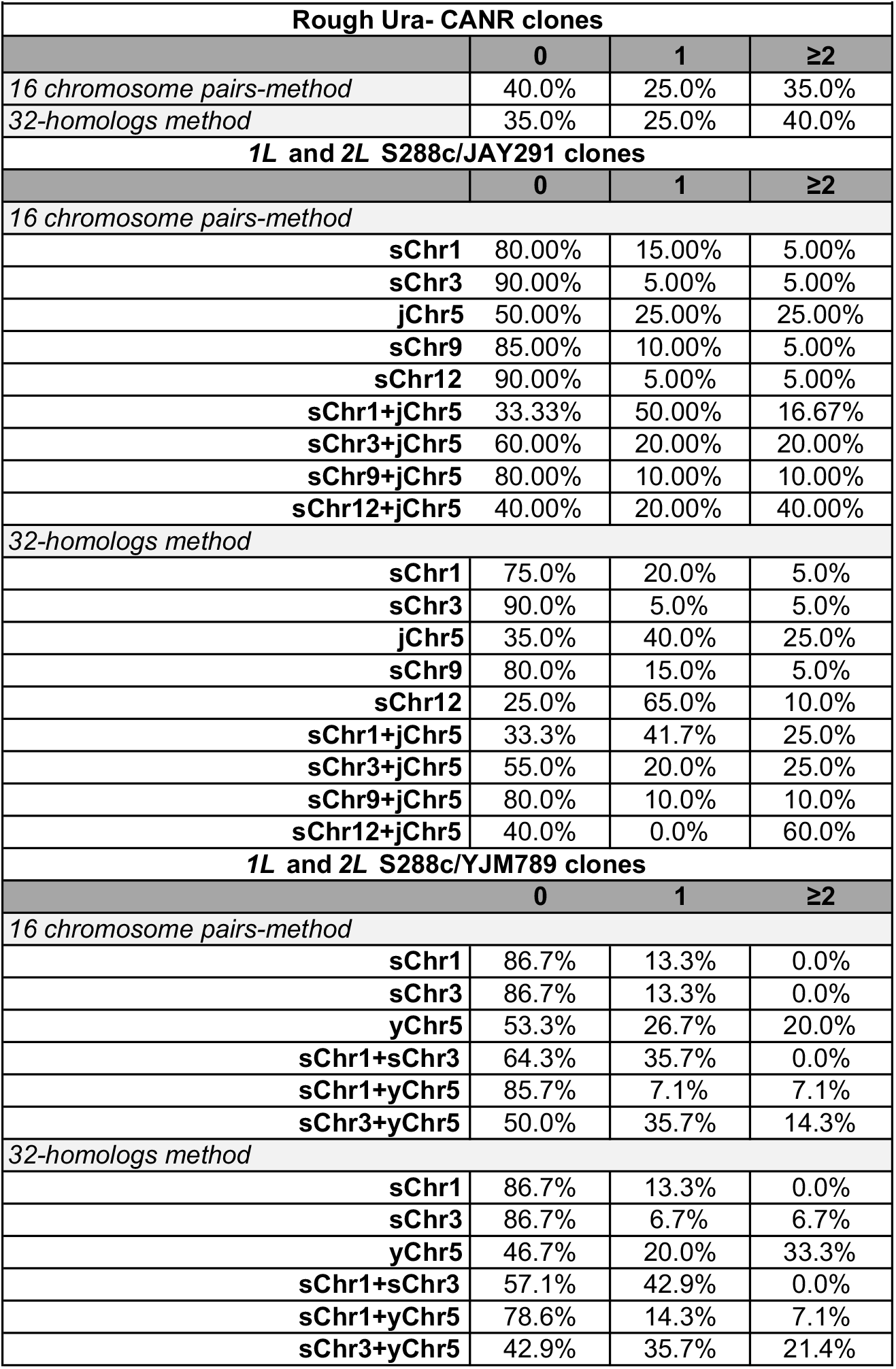
Analysis of the frequency of sequenced clones possessing unselected CCNAs.

**Supplementary Table 7.**
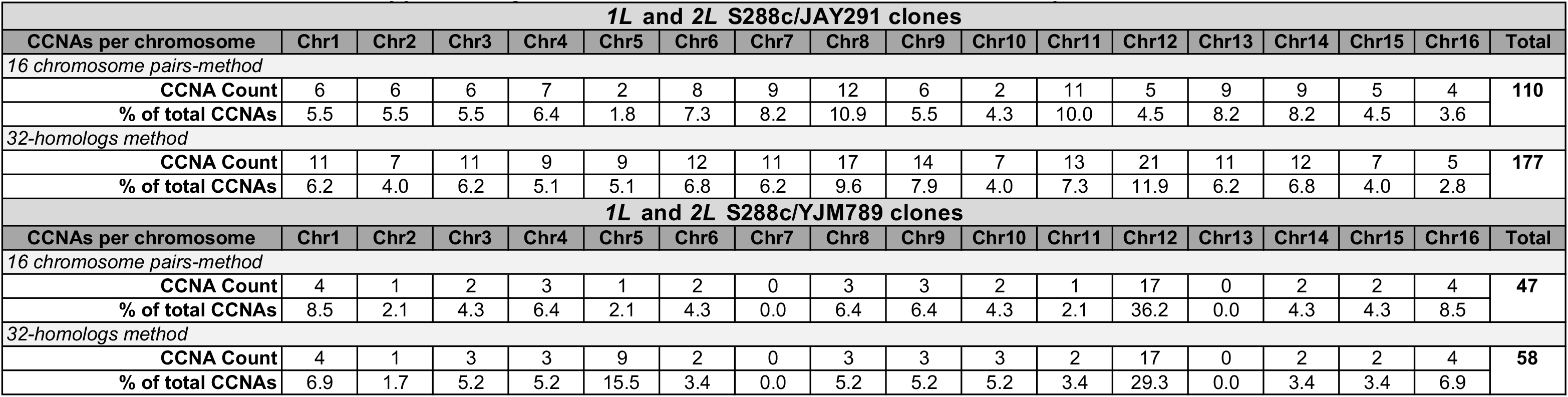
Distribution of unselected CCNAs sorted by chromosome.

**Supplementary Table 8.**
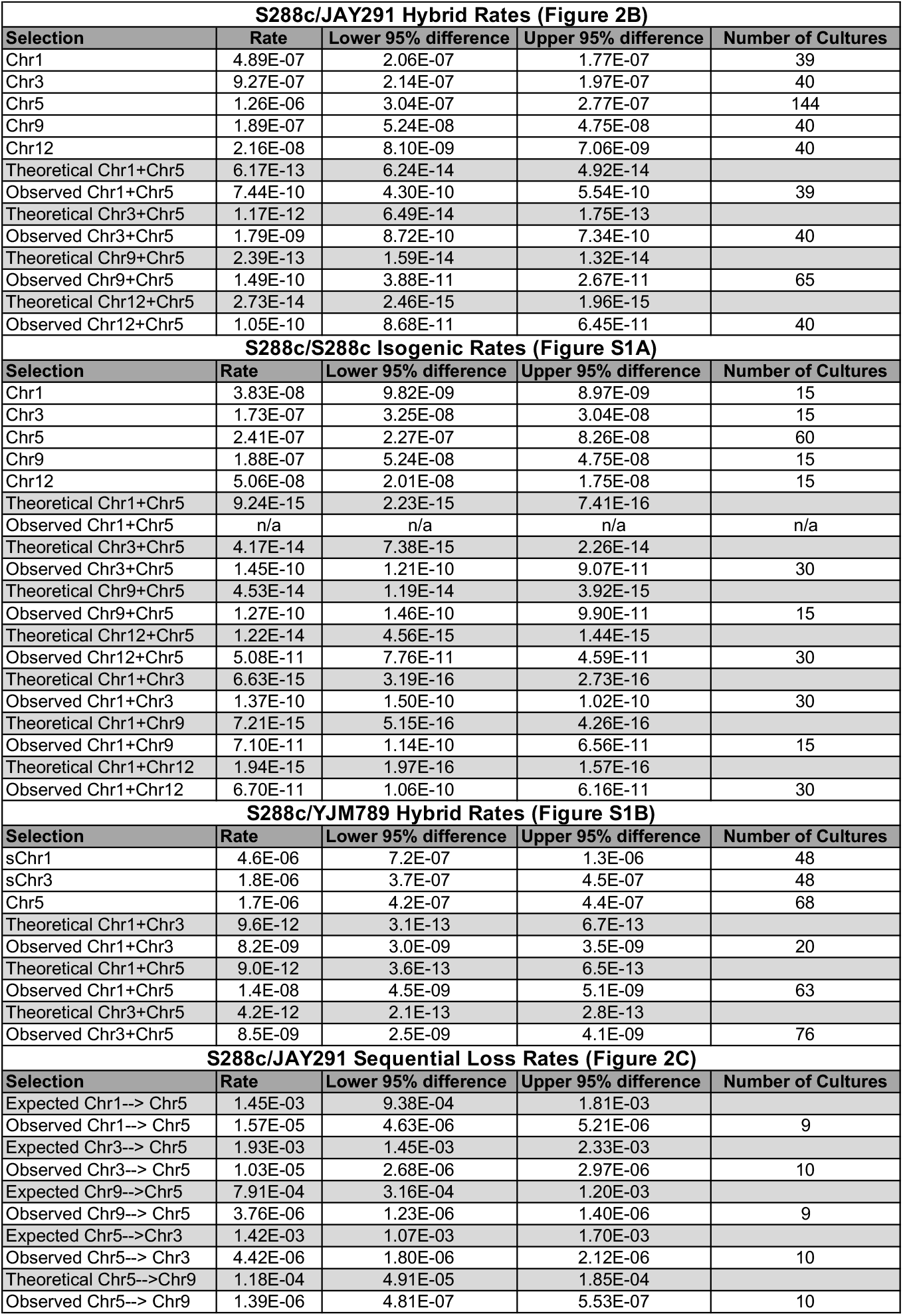
Rates of 1L and 2L chromosome loss calculated using fluctuation analysis. Light grey rows denote Theoretical 2L rate predictions calculated from empirically-derived 1L rates for each chromosome in the denoted genetic background.

