## Supplementary Table 1. Yeast strains used in this study. for "Punctuated Aneuploidization of the Budding Yeast Genome"

| **Supplementary Table 1.** Yeast Strains Used in This Study | | | | |
|---|---|---|---|---|
| **Strain** | **Genotype** | **Background** | **Source** | **Description** |
| **S288c/JAY291 Hybrid Experiments** | | | | |
| **JAY297** | *MATa ura3-52 leu2Δ1 trp1Δ63* | S288c | Fred Winston | S288c parent |
| **JAY298** | *MATα ura3-52 leu2Δ1 his3Δ200* | S288c | Fred Winston | S288c parent |
| **JAY1176** | *MATa ura3* | JAY291 | Argueso et al., 2009 | JAY291 parent |
| **JAY2736** | *MATα trp1Δ63 leu2Δ1 can1::NATMX4 ime1::HPHMX* Chr9L(94840):: *CORE3* Chr9R(385386):: *KlURA3* | S288c | LRH | sChr9 |
| **JAY2735** | *MATα trp1Δ63 leu2Δ1 can1::NATMX4 ime1::HPHMX* Chr12L(19747):: *CORE3* Chr12R(402528):: *KlURA3* | S288c | LRH | sChr12 |
| **JAY2777** | *MATα leu2Δ1 can1::NATMX4 ime1::HPHMX* Chr1L(65444 *)::KlURA3* Chr1R(156325 *):: KlURA3* | S288c | LRH | sChr12 |
| **JAY2778** | *MATα can1::NATMX4 ime1::HPHMX Chr3L(91324)::CORE3* Chr3R(155596 *)::KlURA3* | S288c | LRH | sChr3 |
| **JAY2772** | *MATa ura3 ime1::HPHMX hom3::CAN1-KANMX* | JAY291 | LRH | jChr5 |
| **JAY2773** | JAY2736 x JAY2772 | JAY291 x S288c | LRH | sChr9/jChr5 |
| **JAY2775** | JAY2735 x JAY2772 | JAY291 x S288c | LRH | sChr12/jChr5 |
| **JAY2780** | JAY2777 x JAY2772 | JAY291 x S288c | LRH | sChr1/jChr5 |
| **JAY2782** | JAY2778 x JAY2772 | JAY291 x S288c | LRH | sChr3/jChr5 |
| **S288c/S288c Experiments** | | | | |
| **JAY2750** | *MATA ura3Δ52 leu2Δ1 trp1Δ63 ime1::HPHMX hom3::CAN1-KANMX/HOM3* | S288c | LRH | sChr5 |
| **JAY2739** | *MATA leu2Δ1 can1::NATMX4 ime1::HPHMX* Chr1L(65444 *)::KlURA3* Chr1R(156325 *):: KlURA3* | | | |
| **JAY2828** | JAY2777 x JAY2750 | S288c x S288c | LRH | sChr1/sChr5 |
| **JAY2829** | JAY2778 x JAY2750 | S288c x S288c | LRH | sChr3/sChr5 |
| **JAY2830** | JAY2736 x JAY2750 | S288c x S288c | LRH | sChr9/sChr5 |
| **JAY2831** | JAY2735 x JAY2750 | S288c x S288c | LRH | sChr12/sChr5 |
| **JAY2832** | JAY2739 x JAY2778 | S288c x S288c | LRH | sChr1/sChr3 |
| **JAY2833** | JAY2739 x JAY2736 | S288c x S288c | LRH | sChr1/sChr9 |
| **JAY2834** | JAY2739 x JAY2735 | S288c x S288c | LRH | sChr1/sChr12 |
| **S288c/YJM789 Hybrid Experiments** | | | | |
| **JAY308** | *MATα ho::hisG, ura3, gal2* | YJM789 | Pheobe Lee | YJM789 parent |
| **JAY2593** | *MATα ho::hisG, ura3, gal2 ime1::HPHMX trp1::NATMX4 hom3::CAN1-KANMX* | YJM789 | LRH | yChr5 |
| **JAY2632** | *MATA trp1Δ63 can1::NATMX4 ime1::HPHMX* Chr1L(65444 *)::KlURA3* Chr1R(156325 *)::KlURA3* Chr3L(91324 *)CORE3* Chr3R(155596 *)KlURA3* | S288c | LRH | sChr1/yChr3 |
| **JAY2597** | JAY2632 x JAY2593 | S288c x YJM790 | LRH | sChr1/sChr3/yChr5 |
