## Supplementary Table 2. Plasmids used in this study. for "Punctuated Aneuploidization of the Budding Yeast Genome"

| **Supplementary Table 2.** Plasmids used in this study | | |
|---|---|---|
| **Name** | **Details** | **Source** |
| **pJA95** | *klURA3-scURA3-KANMX* | Zhang *et al.* , 2013 |
| **pJA73** | *pFA6a-HPHMX* | Goldstein *et al.,* 1999 |
| **pJA72** | *pFA6a-NATMX* | Goldstein *et al.,* 1999 |
| **pJA94** | *pFA6a-KANMX* | Wach *et al.* , 1994 |
