## Supplementary Table 3. Sequencing and copy number analysis of rough Ura- CANR S288c/JAY291 clones. for "Punctuated Aneuploidization of the Budding Yeast Genome"

**Supplementary Table 3.** Sequencing and copy number analysis of rough Ura- CANR S288c/JAY291 clones. Columns: s=S288c, j=JAY291. Selected CCNAs (jChr5 and sChr12) are highlighted in yellow. Unselected CCNAs are highlighted in red. Note: All CCNAs of sChr12 were UPD-type.

| | | Chr1 | | Chr2 | | Chr3 | | Chr4 | | Chr5 | | Chr6 | | Chr7 | | Chr8 | | Chr9 | | Chr10 | | Chr11 | | Chr12 | | Chr13 | | Chr14 | | Chr15 | | Chr16 | | Other unselected events |
|---|---|---|---|---|---|---|---|---|---|---|---|---|---|---|---|---|---|---|---|---|---|---|---|---|---|---|---|---|---|---|---|---|---|---|
| **Strain** | **Selection** | s | j | s | j | s | j | s | j | s | j | s | j | s | j | s | j | s | j | s | j | s | j | s | j | s | j | s | j | s | j | s | j | |
| LRH260 | rough Ura- CANR | 1 | 1 | 1 | 1 | 1 | 1 | 1 | 1 | 1 | 0 | 1 | 1 | 1 | 1 | 1 | 1 | 1 | 1 | 1 | 1 | 1 | 1 | 0 | 2 | 1 | 1 | 1 | 1 | 1 | 1 | 1 | 1 | |
| LRH262 | rough Ura- CANR | 1 | 1 | 1 | 1 | 1 | 1 | 1 | 1 | 2 | 0 | 1 | 1 | 1 | 1 | 1 | 0 | 1 | 2 | 2 | 0 | 2 | 0 | 0 | 2 | 0 | 1 | 0 | 2 | 1 | 1 | 1 | 1 | |
| LRH263 | rough Ura- CANR | 1 | 1 | 1 | 1 | 1 | 1 | 1 | 1 | 1 | 0 | 1 | 1 | 1 | 1 | 1 | 1 | 1 | 1 | 1 | 1 | 1 | 1 | 0 | 2 | 1 | 1 | 1 | 1 | 1 | 1 | 1 | 1 | |
| LRH264 | rough Ura- CANR | 1 | 1 | 1 | 1 | 1 | 1 | 1 | 1 | 1 | 0 | 1 | 1 | 1 | 1 | 1 | 1 | 1 | 1 | 1 | 1 | 1 | 1 | 0 | 2 | 1 | 1 | 0 | 1 | 1 | 1 | 1 | 1 | LOH JAY291 homozygous; Chr8 from 105958-TEL8R |
| LRH265 | rough Ura- CANR | 1 | 1 | 1 | 1 | 1 | 1 | 1 | 1 | 2 | 0 | 1 | 1 | 1 | 1 | 1 | 1 | 1 | 1 | 1 | 1 | 1 | 1 | 0 | 2 | 1 | 1 | 1 | 1 | 1 | 1 | 1 | 1 | |
| LRH266 | rough Ura- CANR | 1 | 1 | 2 | 0 | 2 | 1 | 1 | 1 | 2 | 0 | 2 | 1 | 1 | 1 | 2 | 1 | 1 | 1 | 1 | 1 | 1 | 1 | 0 | 2 | 1 | 1 | 1 | 1 | 1 | 1 | 1 | 1 | |
| LRH267 | rough Ura- CANR | 1 | 1 | 1 | 1 | 2 | 1 | 1 | 1 | 1 | 0 | 2 | 1 | 1 | 1 | 1 | 1 | 1 | 1 | 1 | 1 | 1 | 1 | 0 | 2 | 1 | 1 | 1 | 1 | 1 | 2 | 1 | 1 | |
| LRH268 | rough Ura- CANR | 1 | 1 | 1 | 1 | 1 | 1 | 1 | 1 | 1 | 0 | 1 | 1 | 1 | 1 | 1 | 1 | 1 | 1 | 1 | 1 | 1 | 1 | 0 | 2 | 1 | 1 | 1 | 1 | 1 | 1 | 1 | 1 | |
| LRH269 | rough Ura- CANR | 1 | 1 | 1 | 1 | 1 | 1 | 1 | 1 | 1 | 0 | 1 | 1 | 1 | 1 | 1 | 1 | 1 | 1 | 1 | 1 | 1 | 1 | 0 | 2 | 1 | 1 | 1 | 1 | 1 | 1 | 1 | 1 | |
| LRH270 | rough Ura- CANR | 1 | 1 | 1 | 1 | 1 | 1 | 1 | 1 | 1 | 0 | 1 | 1 | 1 | 1 | 1 | 1 | 1 | 1 | 1 | 1 | 1 | 0 | 0 | 2 | 1 | 1 | 1 | 1 | 1 | 1 | 1 | 1 | |
| LRH271 | rough Ura- CANR | 2 | 2 | 0 | 2 | 2 | 2 | 2 | 0 | 2 | 0 | 0 | 2 | 0 | 2 | 0 | 2 | 2 | 2 | 0 | 2 | 2 | 0 | 0 | 2 | 2 | 0 | 2 | 0 | 2 | 0 | 2 | 0 | |
| LRH272 | rough Ura- CANR | 1 | 1 | 1 | 1 | 1 | 1 | 1 | 1 | 1 | 0 | 1 | 1 | 1 | 1 | 1 | 1 | 1 | 1 | 1 | 1 | 1 | 1 | 0 | 2 | 1 | 1 | 1 | 0 | 1 | 1 | 1 | 1 | |
| LRH273 | rough Ura- CANR | 1 | 1 | 1 | 1 | 2 | 1 | 1 | 1 | 2 | 0 | 1 | 1 | 1 | 1 | 1 | 1 | 1 | 1 | 0 | 2 | 1 | 1 | 0 | 2 | 0 | 2 | 1 | 1 | 1 | 1 | 1 | 1 | |
| LRH274 | rough Ura- CANR | 1 | 1 | 1 | 1 | 1 | 1 | 1 | 1 | 1 | 0 | 1 | 1 | 1 | 1 | 1 | 1 | 1 | 1 | 1 | 1 | 1 | 1 | 0 | 2 | 1 | 1 | 1 | 1 | 1 | 1 | 1 | 1 | |
| LRH275 | rough Ura- CANR | 1 | 1 | 1 | 1 | 1 | 1 | 1 | 1 | 1 | 0 | 1 | 1 | 1 | 1 | 1 | 1 | 1 | 1 | 1 | 1 | 1 | 1 | 0 | 2 | 1 | 1 | 1 | 1 | 1 | 1 | 1 | 1 | |
| LRH276 | rough Ura- CANR | 1 | 1 | 1 | 1 | 1 | 1 | 1 | 1 | 1 | 0 | 1 | 1 | 1 | 1 | 1 | 1 | 1 | 1 | 1 | 1 | 1 | 1 | 0 | 2 | 1 | 1 | 1 | 1 | 1 | 1 | 1 | 1 | |
| LRH277 | rough Ura- CANR | 1 | 1 | 1 | 1 | 1 | 1 | 2 | 1 | 2 | 0 | 1 | 2 | 1 | 1 | 1 | 2 | 1 | 2 | 1 | 2 | 1 | 1 | 0 | 2 | 1 | 1 | 1 | 1 | 1 | 1 | 1 | 1 | |
| LRH278 | rough Ura- CANR | 1 | 1 | 1 | 1 | 1 | 1 | 1 | 1 | 1 | 0 | 1 | 1 | 1 | 1 | 1 | 0 | 1 | 1 | 1 | 1 | 1 | 1 | 0 | 2 | 1 | 1 | 1 | 1 | 1 | 1 | 1 | 1 | |
| LRH279 | rough Ura- CANR | 0 | 1 | 0 | 1 | 1 | 1 | 1 | 0 | 2 | 0 | 1 | 1 | 0 | 1 | 1 | 0 | 1 | 1 | 1 | 1 | 1 | 1 | 0 | 2 | 1 | 0 | 0 | 1 | 1 | 1 | 1 | 0 | |
| LRH280 | rough Ura- CANR | 1 | 1 | 2 | 1 | 2 | 1 | 1 | 2 | 2 | 0 | 1 | 2 | 2 | 1 | 1 | 2 | 1 | 2 | 2 | 1 | 2 | 2 | 0 | 2 | 1 | 2 | 1 | 1 | 2 | 2 | 2 | 1 | |
