## Supplementary Table S6. Analysis of the frequency of sequenced clones possessing unselected CCNAs. for "Punctuated Aneuploidization of the Budding Yeast Genome"

**Supplementary Table 6**. Analysis of the frequency of sequenced clones possessing unselected CCNAs.

| | | | |
|---|---|---|---|
| **Rough Ura- CANR clones** | | | |
| | **0** | **1** | **≥2** |
| *16 chromosome pairs-method* | 40.0% | 25.0% | 35.0% |
| *32-homologs method* | 35.0% | 25.0% | 40.0% |
| ***1L* and *2L* S288c/JAY291 clones** | | | |
| | **0** | **1** | **≥2** |
| *16 chromosome pairs-method* | | | |
| **sChr1** | 80.00% | 15.00% | 5.00% |
| **sChr3** | 90.00% | 5.00% | 5.00% |
| **jChr5** | 50.00% | 25.00% | 25.00% |
| **sChr9** | 85.00% | 10.00% | 5.00% |
| **sChr12** | 90.00% | 5.00% | 5.00% |
| **sChr1+jChr5** | 33.33% | 50.00% | 16.67% |
| **sChr3+jChr5** | 60.00% | 20.00% | 20.00% |
| **sChr9+jChr5** | 80.00% | 10.00% | 10.00% |
| **sChr12+jChr5** | 40.00% | 20.00% | 40.00% |
| *32-homologs method* | | | |
| **sChr1** | 75.0% | 20.0% | 5.0% |
| **sChr3** | 90.0% | 5.0% | 5.0% |
| **jChr5** | 35.0% | 40.0% | 25.0% |
| **sChr9** | 80.0% | 15.0% | 5.0% |
| **sChr12** | 25.0% | 65.0% | 10.0% |
| **sChr1+jChr5** | 33.3% | 41.7% | 25.0% |
| **sChr3+jChr5** | 55.0% | 20.0% | 25.0% |
| **sChr9+jChr5** | 80.0% | 10.0% | 10.0% |
| **sChr12+jChr5** | 40.0% | 0.0% | 60.0% |
| ***1L* and *2L* S288c/YJM789 clones** | | | |
| | **0** | **1** | **≥2** |
| *16 chromosome pairs-method* | | | |
| **sChr1** | 86.7% | 13.3% | 0.0% |
| **sChr3** | 86.7% | 13.3% | 0.0% |
| **yChr5** | 53.3% | 26.7% | 20.0% |
| **sChr1+sChr3** | 64.3% | 35.7% | 0.0% |
| **sChr1+yChr5** | 85.7% | 7.1% | 7.1% |
| **sChr3+yChr5** | 50.0% | 35.7% | 14.3% |
| *32-homologs method* | | | |
| **sChr1** | 86.7% | 13.3% | 0.0% |
| **sChr3** | 86.7% | 6.7% | 6.7% |
| **yChr5** | 46.7% | 20.0% | 33.3% |
| **sChr1+sChr3** | 57.1% | 42.9% | 0.0% |
| **sChr1+yChr5** | 78.6% | 14.3% | 7.1% |
| **sChr3+yChr5** | 42.9% | 35.7% | 21.4% |
