## Supplementary Table S7. Proportion of unselected CCNAs affecting each chromosome. for "Punctuated Aneuploidization of the Budding Yeast Genome"

**Supplementary Table 7.** Distribution of unselected CCNAs sorted by chromosome.

| ***1L* and *2L* S288c/JAY291 clones** | | | | | | | | | | | | | | | | | |
|---|---|---|---|---|---|---|---|---|---|---|---|---|---|---|---|---|---|
| **CCNAs per chromosome** | **Chr1** | **Chr2** | **Chr3** | **Chr4** | **Chr5** | **Chr6** | **Chr7** | **Chr8** | **Chr9** | **Chr10** | **Chr11** | **Chr12** | **Chr13** | **Chr14** | **Chr15** | **Chr16** | **Total** |
| *16 chromosome pairs-method* | | | | | | | | | | | | | | | | | |
| **CCNA Count** | 6 | 6 | 6 | 7 | 2 | 8 | 9 | 12 | 6 | 2 | 11 | 5 | 9 | 9 | 5 | 4 | **110** |
| **% of total CCNAs** | 5.5 | 5.5 | 5.5 | 6.4 | 1.8 | 7.3 | 8.2 | 10.9 | 5.5 | 4.3 | 10.0 | 4.5 | 8.2 | 8.2 | 4.5 | 3.6 | |
| *32-homologs method* | | | | | | | | | | | | | | | | | |
| **CCNA Count** | 11 | 7 | 11 | 9 | 9 | 12 | 11 | 17 | 14 | 7 | 13 | 21 | 11 | 12 | 7 | 5 | **177** |
| **% of total CCNAs** | 6.2 | 4.0 | 6.2 | 5.1 | 5.1 | 6.8 | 6.2 | 9.6 | 7.9 | 4.0 | 7.3 | 11.9 | 6.2 | 6.8 | 4.0 | 2.8 | |
| ***1L* and *2L* S288c/YJM789 clones** | | | | | | | | | | | | | | | | | |
| **CCNAs per chromosome** | **Chr1** | **Chr2** | **Chr3** | **Chr4** | **Chr5** | **Chr6** | **Chr7** | **Chr8** | **Chr9** | **Chr10** | **Chr11** | **Chr12** | **Chr13** | **Chr14** | **Chr15** | **Chr16** | **Total** |
| *16 chromosome pairs-method* | | | | | | | | | | | | | | | | | |
| **CCNA Count** | 4 | 1 | 2 | 3 | 1 | 2 | 0 | 3 | 3 | 2 | 1 | 17 | 0 | 2 | 2 | 4 | **47** |
| **% of total CCNAs** | 8.5 | 2.1 | 4.3 | 6.4 | 2.1 | 4.3 | 0.0 | 6.4 | 6.4 | 4.3 | 2.1 | 36.2 | 0.0 | 4.3 | 4.3 | 8.5 | |
| *32-homologs method* | | | | | | | | | | | | | | | | | |
| **CCNA Count** | 4 | 1 | 3 | 3 | 9 | 2 | 0 | 3 | 3 | 3 | 2 | 17 | 0 | 2 | 2 | 4 | **58** |
| **% of total CCNAs** | 6.9 | 1.7 | 5.2 | 5.2 | 15.5 | 3.4 | 0.0 | 5.2 | 5.2 | 5.2 | 3.4 | 29.3 | 0.0 | 3.4 | 3.4 | 6.9 | |
