## Supplementary Table S8. Rates of 1L and 2L chromosome loss calculated using fluctuation analysis. for "Punctuated Aneuploidization of the Budding Yeast Genome"

Supplementary Table 8. Rates of 1L and 2L chromosome loss calculated using fluctuation analysis. Light grey rows denote Theoretical 2L rate predictions calculated from empirically-derived 1L rates for each chromosome in the denoted genetic background.

| **S288c/JAY291 Hybrid Rates (Figure 2B)** | | | | |
|---|---|---|---|---|
| **Selection** | **Rate** | **Lower 95% difference** | **Upper 95% difference** | **Number of Cultures** |
| Chr1 | 4.89E-07 | 2.06E-07 | 1.77E-07 | 39 |
| Chr3 | 9.27E-07 | 2.14E-07 | 1.97E-07 | 40 |
| Chr5 | 1.26E-06 | 3.04E-07 | 2.77E-07 | 144 |
| Chr9 | 1.89E-07 | 5.24E-08 | 4.75E-08 | 40 |
| Chr12 | 2.16E-08 | 8.10E-09 | 7.06E-09 | 40 |
| Theoretical Chr1+Chr5 | 6.17E-13 | 6.24E-14 | 4.92E-14 | |
| Observed Chr1+Chr5 | 7.44E-10 | 4.30E-10 | 5.54E-10 | 39 |
| Theoretical Chr3+Chr5 | 1.17E-12 | 6.49E-14 | 1.75E-13 | |
| Observed Chr3+Chr5 | 1.79E-09 | 8.72E-10 | 7.34E-10 | 40 |
| Theoretical Chr9+Chr5 | 2.39E-13 | 1.59E-14 | 1.32E-14 | |
| Observed Chr9+Chr5 | 1.49E-10 | 3.88E-11 | 2.67E-11 | 65 |
| Theoretical Chr12+Chr5 | 2.73E-14 | 2.46E-15 | 1.96E-15 | |
| Observed Chr12+Chr5 | 1.05E-10 | 8.68E-11 | 6.45E-11 | 40 |
| **S288c/S288c Isogenic Rates (Figure S1A)** | | | | |
| **Selection** | **Rate** | **Lower 95% difference** | **Upper 95% difference** | **Number of Cultures** |
| Chr1 | 3.83E-08 | 9.82E-09 | 8.97E-09 | 15 |
| Chr3 | 1.73E-07 | 3.25E-08 | 3.04E-08 | 15 |
| Chr5 | 2.41E-07 | 2.27E-07 | 8.26E-08 | 60 |
| Chr9 | 1.88E-07 | 5.24E-08 | 4.75E-08 | 15 |
| Chr12 | 5.06E-08 | 2.01E-08 | 1.75E-08 | 15 |
| Theoretical Chr1+Chr5 | 9.24E-15 | 2.23E-15 | 7.41E-16 | |
| Observed Chr1+Chr5 | n/a | n/a | n/a | n/a |
| Theoretical Chr3+Chr5 | 4.17E-14 | 7.38E-15 | 2.26E-14 | |
| Observed Chr3+Chr5 | 1.45E-10 | 1.21E-10 | 9.07E-11 | 30 |
| Theoretical Chr9+Chr5 | 4.53E-14 | 1.19E-14 | 3.92E-15 | |
| Observed Chr9+Chr5 | 1.27E-10 | 1.46E-10 | 9.90E-11 | 15 |
| Theoretical Chr12+Chr5 | 1.22E-14 | 4.56E-15 | 1.44E-15 | |
| Observed Chr12+Chr5 | 5.08E-11 | 7.76E-11 | 4.59E-11 | 30 |
| Theoretical Chr1+Chr3 | 6.63E-15 | 3.19E-16 | 2.73E-16 | |
| Observed Chr1+Chr3 | 1.37E-10 | 1.50E-10 | 1.02E-10 | 30 |
| Theoretical Chr1+Chr9 | 7.21E-15 | 5.15E-16 | 4.26E-16 | |
| Observed Chr1+Chr9 | 7.10E-11 | 1.14E-10 | 6.56E-11 | 15 |
| Theoretical Chr1+Chr12 | 1.94E-15 | 1.97E-16 | 1.57E-16 | |
| Observed Chr1+Chr12 | 6.70E-11 | 1.06E-10 | 6.16E-11 | 30 |
| **S288c/YJM789 Hybrid Rates (Figure S1B)** | | | | |
| **Selection** | **Rate** | **Lower 95% difference** | **Upper 95% difference** | **Number of Cultures** |
| sChr1 | 4.6E-06 | 7.2E-07 | 1.3E-06 | 48 |
| sChr3 | 1.8E-06 | 3.7E-07 | 4.5E-07 | 48 |
| Chr5 | 1.7E-06 | 4.2E-07 | 4.4E-07 | 68 |
| Theoretical Chr1+Chr3 | 9.6E-12 | 3.1E-13 | 6.7E-13 | |
| Observed Chr1+Chr3 | 8.2E-09 | 3.0E-09 | 3.5E-09 | 20 |
| Theoretical Chr1+Chr5 | 9.0E-12 | 3.6E-13 | 6.5E-13 | |
| Observed Chr1+Chr5 | 1.4E-08 | 4.5E-09 | 5.1E-09 | 63 |
| Theoretical Chr3+Chr5 | 4.2E-12 | 2.1E-13 | 2.8E-13 | |
| Observed Chr3+Chr5 | 8.5E-09 | 2.5E-09 | 4.1E-09 | 76 |
| **S288c/JAY291 Sequential Loss Rates (Figure 2C)** | | | | |
| **Selection** | **Rate** | **Lower 95% difference** | **Upper 95% difference** | **Number of Cultures** |
| Expected Chr1--> Chr5 | 1.45E-03 | 9.38E-04 | 1.81E-03 | |
| Observed Chr1--> Chr5 | 1.57E-05 | 4.63E-06 | 5.21E-06 | 9 |
| Expected Chr3--> Chr5 | 1.93E-03 | 1.45E-03 | 2.33E-03 | |
| Observed Chr3--> Chr5 | 1.03E-05 | 2.68E-06 | 2.97E-06 | 10 |
| Expected Chr9-->Chr5 | 7.91E-04 | 3.16E-04 | 1.20E-03 | |
| Observed Chr9--> Chr5 | 3.76E-06 | 1.23E-06 | 1.40E-06 | 9 |
| Expected Chr5-->Chr3 | 1.42E-03 | 1.07E-03 | 1.70E-03 | |
| Observed Chr5--> Chr3 | 4.42E-06 | 1.80E-06 | 2.12E-06 | 10 |
| Theoretical Chr5-->Chr9 | 1.18E-04 | 4.91E-05 | 1.85E-04 | |
| Observed Chr5--> Chr9 | 1.39E-06 | 4.81E-07 | 5.53E-07 | 10 |
